# Prenatal maternal stress is associated with increased sensitivity to neuropathic pain and sex-specific epigenetic and transcriptomic dynamics in the prefrontal cortex

**DOI:** 10.1101/2025.11.27.691002

**Authors:** Arthur Vovk, Stéphanie Grégoire, Zofia Barel, Laura S Stone, Elad Lax

## Abstract

Prenatal maternal stress (PNS) is a common early-life adversity. PNS has been linked to greater vulnerability to chronic pain in the offspring. PNS results in increased hypersensitivity after Chronic Constriction Injury (CCI) of the sciatic nerve, a common rodent model for chronic neuropathic pain. These behavioral effects are accompanied by altered levels of enzymes that regulate DNA methylation in the frontal cortex. DNA methylation, the addition of a methyl group onto cytosine bases of cytosine-guanine (CpG) sites in the genome, is an epigenetic mechanism by which life experiences can reprogram gene expression. The goal of this study was to identify differentially methylated regions that might contribute to the increased pain sensitivity following nerve injury in adulthood. Prior studies indicate widespread, sex-specific transcriptomic changes and numerous differentially methylated regions (DMRs) in the frontal cortex following PNS and nerve injury. Here we performed genome wide DNA methylation and transcriptome-wide RNAseq analysis in the frontal cortex of male and female mice that underwent PNS, CCI or both. Our analysis revealed sex-dependent changes in DNA methylation and mRNA expression, including in pathways related to neuronal and brain development, axonogenesis and synaptic regulation. The effect of DNA methylation on mRNA expression was tested in a subset of target genes. These studies suggest a role for DNA methylation in embedding increased risk for chronic pain in adulthood associated with early-life adversity.

## Introduction

Chronic pain is one of the most prevalent health problems worldwide and a leading cause of disability, with major implications for public health and health-care systems. In the United States, it was estimated that over 50 million adults (∼20%) experience chronic pain, and more than 20 million live with high-impact pain that substantially limits daily activities^1,2^. Beyond physical suffering, chronic pain has been shown to cause cognitive disturbances, implicating changes within the central nervous system that promote emotion regulation and executive control^3,4^. Neuroimaging demonstrates that disrupted corticostriatal connectivity predicts transition from acute to chronic back pain^5^, and that clinical treatment can partially reverse abnormal brain anatomy and function, including in prefrontal regions^6^. In parallel, longitudinal animal work shows structural and functional network alterations with neuropathic pain, with distinct changes persisting in a chronic pain state, reinforcing the concept of pain as a systems-level disorder of neuroplasticity^7^.

Prenatal maternal stress (PNS) is the maternal exposure to psychological and/or physical stress during pregnancy. It has emerged as a critical environmental factor that shapes offspring neurodevelopment, causing long-term effects^8^. Exposure of humans to PNS demonstrates that prenatal stress produces stable DNA methylation signatures in the offspring^9^. Similarly, prenatal maternal depression has been linked to epigenetic alterations in the offspring, suggesting an in-utero program of stress-responsive pathways^10^.

Evidence across multiple studies indicates that early stress exposure increases later pain vulnerability^11^. In the 1958 British Birth Cohort, childhood adversity predicted chronic widespread pain in adulthood^12^. Young adults with childhood adversity show a higher likelihood of chronic pain conditions^13^, and populations with childhood abuse exhibit higher rates of co-occurring depression and chronic pain^14^. These observations indicate that prenatal and early-life stress can increase the risk for pain and pain sensitivity in later life.

Preclinical studies show that developmental stress alters neuroplasticity, resulting in lasting behavioral and cognitive effects^15^. Early-life stress enhances adult nerve injury-induced hypersensitivity in both mechanical and thermal tests^16^. Most directly, we demonstrated that PNS sensitizes offspring to neuropathic pain following chronic constriction injury (CCI), accompanied by sex-specific changes in expression of epigenetic- and stress-related genes^17^. Together, these findings support a model in which PNS establishes a latent vulnerability that emerges following later-life injury.

Long-lasting effects of PNS are widely attributed to epigenetic regulation, particularly DNA methylation, which can modulate transcription without altering the DNA sequence^18^. Studies have shown that early-life experiences can alter stress-related gene transcription through DNA methylation. In rodents, variation in maternal care induces methylation changes at the glucocorticoid receptor (NR3C1), leading to stable differences in HPA axis responsivity the persist into adulthood^19^. In humans, childhood abuse is associated with hypermethylation of NR3C1 in the hippocampus, linking adversity to lasting stress dysregulation^20^.

Chronic pain causes cortical epigenomic remodeling. After peripheral nerve injury, the prefrontal cortex (PFC) exhibits reversible global methylation changes^21^ and transcriptome-wide remodeling^22^. Importantly, shared pain-related methylation signatures appear in both PFC and peripheral T cells, suggesting systemic coordination and translational biomarker potential^23^.

Plasticity-relevant genes are under epigenetic control in pain contexts. Demethylation of BDNF in dorsal root ganglion neurons contributes to opioid-induced hypersensitivity^24^. Activity-dependent transcription factors (MEF2, Npas4) linked to excitatory–inhibitory balance are also epigenetically regulated, showing a connection between methylation dynamics to synaptic remodeling^25,26^. Epigenetic enzymes are key to this control: Ten-Eleven Translocation (TET1-3) family of dioxygenases oxidize methylated cytosines, and DNA Methyltransferases DNMT1, DNMT3A and DNMT3B) catalyze DNA methylation contributes to the etiology of chronic pain and mediates chronic pain-related depressive like behaviors^27–31^. The frontal cortex, especially the medial prefrontal cortex (mPFC), integrates sensory, affective, and cognitive components of pain within cortico-limbic circuits^4^. In humans, abnormal mPFC connectivity is associated with the development of chronic pain^5^, and surgical or behavioral treatment can normalize prefrontal function^6^. In rodents, neuropathic pain induces remodeling of mPFC circuits that drive anxiety-like behaviors and cognitive deficits^32^, with vmPFC and dmPFC projection neurons shaping both affective and nociceptive outputs^33,34^. Long-term cortical synaptic plasticity is increasingly recognized as a driver of both persistent pain and emotional dysregulation^35^.

Crucially, these circuit changes are stabilized by epigenetic mechanisms. Following nerve injury, the frontal cortex undergoes long-lasting methylation reprogramming detectable up to one year post-injury^36^. Treatment with the methyl donor S-adenosylmethionine reverses pain-linked DNA methylation changes and reduces mechanical hypersensitivity, showing these epigenetic changes are reversible^37^. Overall, PNS shifts cortical methylation and demethylation enzymes^17^, priming pain circuits for stronger responses to injury and leading to heightened pain sensitivity.

Despite these insights, a critical knowledge gap remains: Does PNS reprogram DNA methylation in the frontal cortex such that transcriptional responses following nerve injury in adulthood lead to increased pain susceptibility? In addition, since most previous studies included only male mice, the potential sex-specific effects of PNS on chronic pain, DNA methylation, and gene-expression is poorly explored. Addressing these gaps will provide insights into the developmental origins of pain sensitivity and help achieve targeted, epigenetically guided interventions. To address this, we collected and sequenced DNA and RNA samples from the PFC of male and female adult offspring of PNS and control mouse dams who underwent Chronic Constriction Injury (CCI) and sham control surgery. The effects of both PNS and CCI on frontal cortex genome-wide DNA methylation and gene-expression patterns were explored.

## Methods

All animal procedures and sample collection, including RNA and DNA extraction, were performed at the Lab of Prof. Laura Stone at the Alan Edwards Centre for Research on Pain, McGill University, Montreal, QC, Canada, and the Department of Anesthesiology, Medical School, the University of Minnesota, MN, USA. The samples used for the analyses described here were obtained from the same experimental animals and prepared as previously described^17^.

### Animals

Pregnant CD1 females (n = 6) were obtained on embryonic day (E) 9 from Charles River Laboratories (St-Constant, QC, Canada). Each dam was housed individually under a 12-h light/dark cycle in a temperature-controlled facility, using ventilated polycarbonate cages (Allentown, NJ, USA) with corncob bedding (Teklad 7097, Envigo, UK) and cotton nesting material for enrichment. Food (2092X Global Soy Protein-Free Extruded Rodent Diet, irradiated) and water were available ad libitum.

All animal procedures were reviewed and approved by the McGill University Animal Care Committee and carried out in accordance with the guidelines of the Canadian Council on Animal Care as well as the International Association for the Study of Pain’s Committee for Research and Ethical Issues^38,39^.

### Prenatal Stress Procedure

From E13 to E17, a subset of pregnant mice (n = 3) underwent a repeated prenatal stress protocol, whereas controls (n = 3) remained undisturbed in their home cages. Stress exposure consisted of restraining the dam in a transparent 500-ml glass cylinder (4.5 cm diameter, >30 cm length) partially filled with 5 mm of water, under bright illumination. Each dam experienced three sessions per day (08:00–10:00 h, 12:00–14:00 h, and 16:00–18:00 h), lasting 45 min per session, as adapted from a previously described paradigm^40^.

Following parturition, litters were reared by their biological mothers until weaning, after which offspring were housed in same-sex groups of two to four under standard laboratory conditions. At three months of age, animals were subjected to baseline behavioral testing before undergoing either chronic constriction injury (CCI) of the sciatic nerve or sham surgery (n = 9–11 per group). Post-surgical assessments of mechanical and cold sensitivity were performed at 1, 2, 3, and 6 weeks, followed by evaluation of anxiety- and depressive-like behaviors at 6–7 weeks post-injury.

At the conclusion of behavioral testing, mice were anesthetized with isoflurane and sacrificed by decapitation. Brains were rapidly extracted and dissected on ice according to stereotaxic coordinates (Paxinos & Franklin, 2004). Tissue punches corresponding to the frontal cortex (+1 to +2.5 mm anterior to bregma, −0.75 to +0.75 mm lateral, −2 to −3.5 mm ventral) and hippocampus (−1 to −3.5 mm anterior, −3.5 to +3.5 mm lateral, −1 to −3.5 mm ventral) were collected, flash-frozen on dry ice, and stored at −80 °C until processing. For all analyses, hemispheres were pooled.

### Chronic Constriction Injury (CCI)

Neuropathic pain was induced using a modified version of the Bennett and Xie method^41^. Under isoflurane anesthesia, the left sciatic nerve was exposed at the mid-thigh level and loosely ligated with four 4-0 chromic gut sutures, spaced approximately 1 mm apart. The overlying muscle and skin were closed with a single 3-0 sterile silk suture, and a topical antibiotic was applied. Sham-operated animals underwent the same exposure procedure without ligation of the nerve. All surgical interventions were performed by the same experimenter to minimize variability.

### Behavioral Assessment of Pain Sensitivity

To assess pain sensitivity in this model, mechanical and cold responses were evaluated in three-month-old offspring at baseline and at 1, 2, 3, and 6 weeks following CCI or sham surgery. After habituation to the testing environment, mechanical sensitivity was measured using calibrated von Frey filaments (0.04–4.0 g; Stoelting Co., IL) applied to the plantar surface of the hind paw, with withdrawal thresholds determined by the up-down method^42^. Cold sensitivity was assessed using a modified acetone drop test, in which 25 μL of acetone was applied to the plantar surface, and the duration of nocifensive behaviors (flinching, licking, biting) was recorded over 1 min^43^.

### RNA and DNA Extraction

Total RNA and DNA was extracted from frontal cortex tissue using the AllPrep DNA/RNA Mini Kit (Qiagen, Hilden, Germany) in accordance with the manufacturer’s instructions.

### RNA Sequencing

Total RNA sample concentrations were quantified with a RiboGreen RNA quantification kit. RNA integrity was assessed using Tapestation. Poly-A selection and sequencing libraries were made at the University of Minnesota Genomic Center. Illumina TruSeq Stranded mRNA reagents were used, and libraries were sequenced on an Illumina NovaSeq platform using 150 bp paired-end reads, following the manufacturer’s recommended protocol with ∼20M reads per sample. Sequencing reads QC was assessed using FASTQC and reads were deduplicated and aligned to the mouse reference genome (mm10) with STAR aligner^44^. Differential gene expression was assessed with DESeq2, applying a false discovery rate (FDR) threshold of 0.05^45^.

### Reverse Transcription and Quantitative Real-Time PCR (qPCR)

For quantitative Real-Time PCR (qPCR), RNA concentration and purity were assessed, and 200 ng of total RNA from each sample was reverse transcribed into cDNA using the High-Capacity cDNA Reverse Transcription Kit (Applied Biosystems, Foster City, USA). Quantitative PCR was performed on a Quant studio 3 (Applied Biosystems) using SsoAdvanced™ Universal SYBR® Green Supermix (Bio-Rad, USA). Each reaction was carried out in triplicate for each sample. Relative gene expression was determined using the ΔΔCt method, with the Control-Sham group serving as the reference. Expression values were normalized to the geometric mean of β-actin Ct values, which were used as endogenous controls.

### Reduced-Representation Bisulfite Sequencing (RRBS)

Reduced-Representation Bisulfite Sequencing (RRBS) was performed to analyze DNA methylation changes in the frontal cortex following maternal prenatal stress and chronic pain exposure. This method provides genome-wide DNA methylation profiles by enriching CpG-rich regions using the restriction enzymes MspI and TaqαI. Genomic DNA was digested with these enzymes and then was subjected to bisulfite conversion using the Zymo-Seq RRBS Library Kit and following the manufacturer’s protocol workflow (Zymo Research).

Sequencing reads QC was assessed using FASTQC, and reads were deduplicated and aligned to the mouse reference genome (mm10) with BISMARK^46^. Differential methylation analysis was conducted to detect differentially methylated regions (DMRs) across experimental groups using the R script MethylKit^47^.

To assess the relationship between DNA methylation and gene expression, we screened the data for overlapping DMRs who exhibit significant differential methylation in more than one RRBS pair-wise comparison and assessed their mRNA expression levels using qPCR.

### Statistical Analysis

Statistical analyses of behavioral data and qPCR was done using two-way ANOVAs followed by post-hoc tests. For RRBS data, pair-wise differential methylation was analyzed using the mann-whitney u test. For RNA-seq analysis the raw read counts were modeled using a Negative Binomial Generalized Linear Model (GLM). The full factorial design model included the two main factors (PNS and CCI) and their interaction term. For the RNA-seq analysis, differential gene expression was determined by modelling the raw read counts using a Generalized Linear Model (GLM) framework. The GLM was fitted using a negative binomial distribution. The model was based on a full factorial design and included all experimental factors: the two main factors, PNS (Prenatal Stress) and CCI (Chronic Constriction Injury), as well as their interaction term (PNS X CCI) to specifically assess if the effect of one factor is dependent on the level of the other. Multiple comparisons were corrected using the Benjamini–Hochberg false discovery rate (adjusted p < 0.05). For DNA methylation, a minimum difference of 5% and ≥10 reads per site was required. For RNA-seq, a minimum of 10 read counts per gene was applied, consistent with DeSeq2 pipeline recommendations^48^. Gene pathway analyses were done with Metascape^49^.

## Results

### Prenatal Stress and Chronic Pain Hypersensitivity

We previously reported the results of a combined prenatal stress and chronic constriction injury (CCI) model. This model shows that PNS results in increased hypersensitivity following nerve injury in the adult mouse offspring alongside anxiety-like behaviors**^17^**Specifically, we demonstrated that offspring from stressed dams exhibit increased mechanical sensitivity following CCI compared to controls (Figure 1)^17^. For the present study, we used brain samples collected from this established cohort to investigate the molecular and epigenetic mechanisms underlying these persistent behavioral alterations.

**Figure 1.**
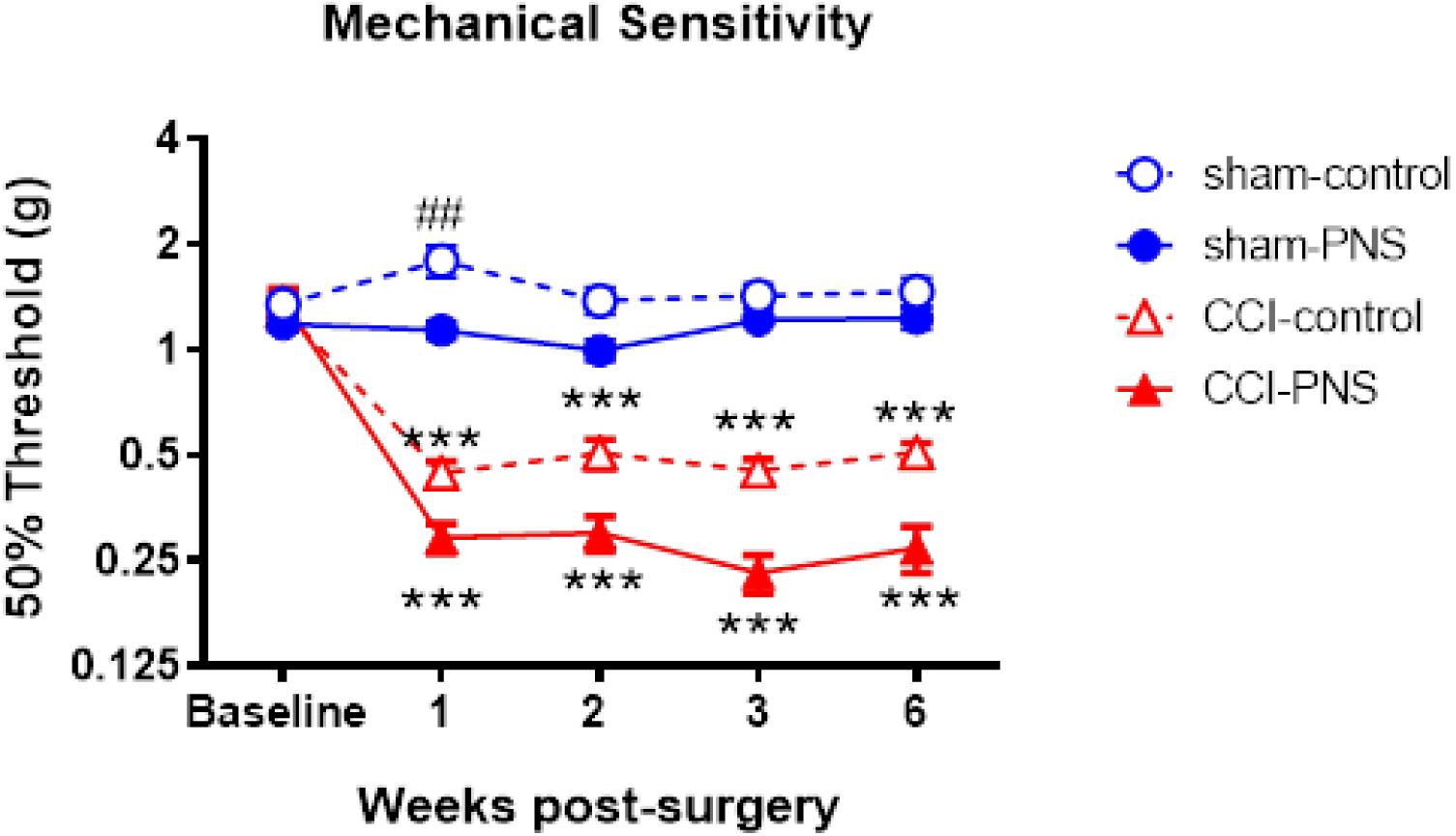
Effect of prenatal maternal stress on pain sensitivity in adult offspring. Mechanical sensitivity thresholds were measured on offspring at baseline and 1, 2, 3, and 6 weeks post-CCI induction in prenatally stressed or control animals. Results are represented with mean ± SEM. 2-way repeated measures ANOVA followed by Tukey multiple comparisons post-hoc test was used. *p < 0.05, **p < 0.01, ***p < 0.001, ##p < 0.01.

### Transcriptomic Alterations Following PNS and CCI

To investigate transcriptional changes associated with PNS and CCI, we performed RNA-seq on frontal cortex samples. Differentially expressed genes (DEGs) were identified for PNS, CCI, and their interaction.

RNA sequencing of frontal cortex tissue identified widespread, sex-dependent, transcriptional alterations associated with prenatal stress, neuropathic injury, and their interaction (Table 1). These results demonstrate that both PNS and CCI drive distinct transcriptional programs, with the interaction capturing additional gene sets not explained by either factor alone. Notably, the magnitude and pattern of regulation differed between sexes, with males showing a particularly strong transcriptional response to CCI, whereas females displayed more extensive dysregulation resulting from the interaction of PNS and CCI.

**Table 1.** Number of differentially expressed genes (DEGs) in the frontal cortex following prenatal stress and chronic constriction injury (CCI). Counts of DEGs in frontal cortex of male and female offspring for each comparison (prenatal stress, CCI, and their interaction). DEGs were defined as genes with adjusted p-value (padj) < 0.05. Interaction refers to genes significantly regulated by the combined effect of prenatal stress and CCI, beyond the main effects of either factor alone.

| Effect | Female (DEGs) | Male (DEGs) |
| --- | --- | --- |
| Prenatal Stress | 369 | 419 |
| CCI (Pain) | 155 | 1086 |
| Interaction | 660 | 22 |

To further visualize these transcriptional changes, volcano plots were generated separately for female and male offspring. In females (Figure 2), PNS (Figure 2A) and CCI (Figure 2B) each induced distinct transcriptional alterations, while the interaction (Figure 2C) produced the broadest transcriptional response. In males (Figure 3), PNS induced relatively less differentially expressed genes (Figure 3A) compared to CCI (Figure 3B), which accounted for the stronger transcriptional shift. The interaction between these two factors showed only a small subset of significantly regulated genes (Figure 3C).

**Figure 2.**
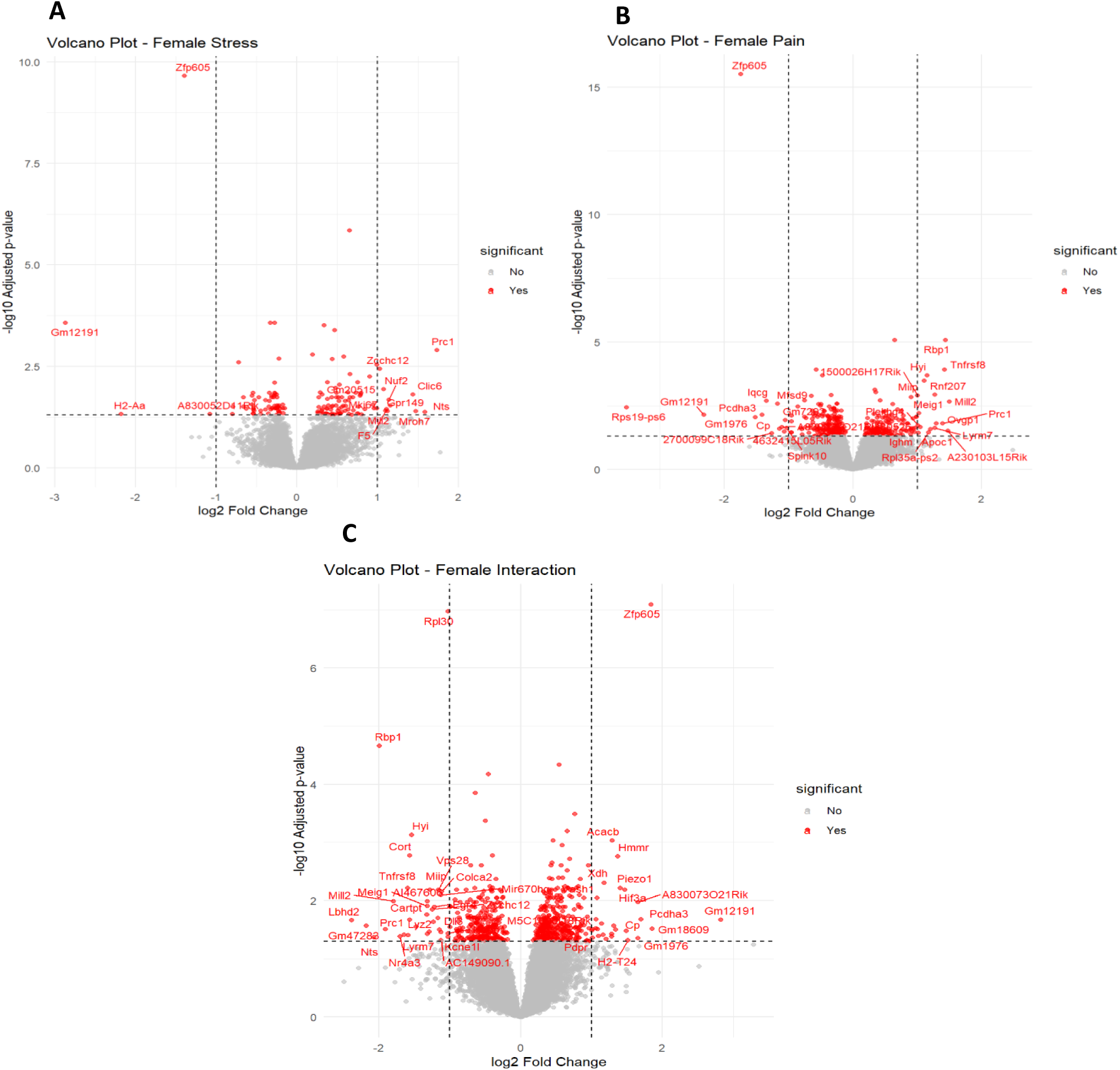
**Differential gene expression in the female frontal cortex following prenatal stress and chronic constriction injury (CCI)**. Differentially expressed genes identified by RNA sequencing. **(A)** Main effect of prenatal stress (prenatal stress vs sham). **(B)** Main effect of neuropathic pain (CCI vs sham). **(C)** Statistical interaction between prenatal stress and CCI, representing genes significantly regulated by the combined effect of both factors beyond their individual contributions. The x-axis represents log₂ fold change, and the y-axis shows the –log₁₀ adjusted *p*-value (FDR < 0.05). Genes passing significance thresholds are highlighted in red, with selected transcripts labeled. This data illustrates distinct and overlapping transcriptional alterations associated with prenatal stress, neuropathic injury, and their interaction.

**Figure 3.**
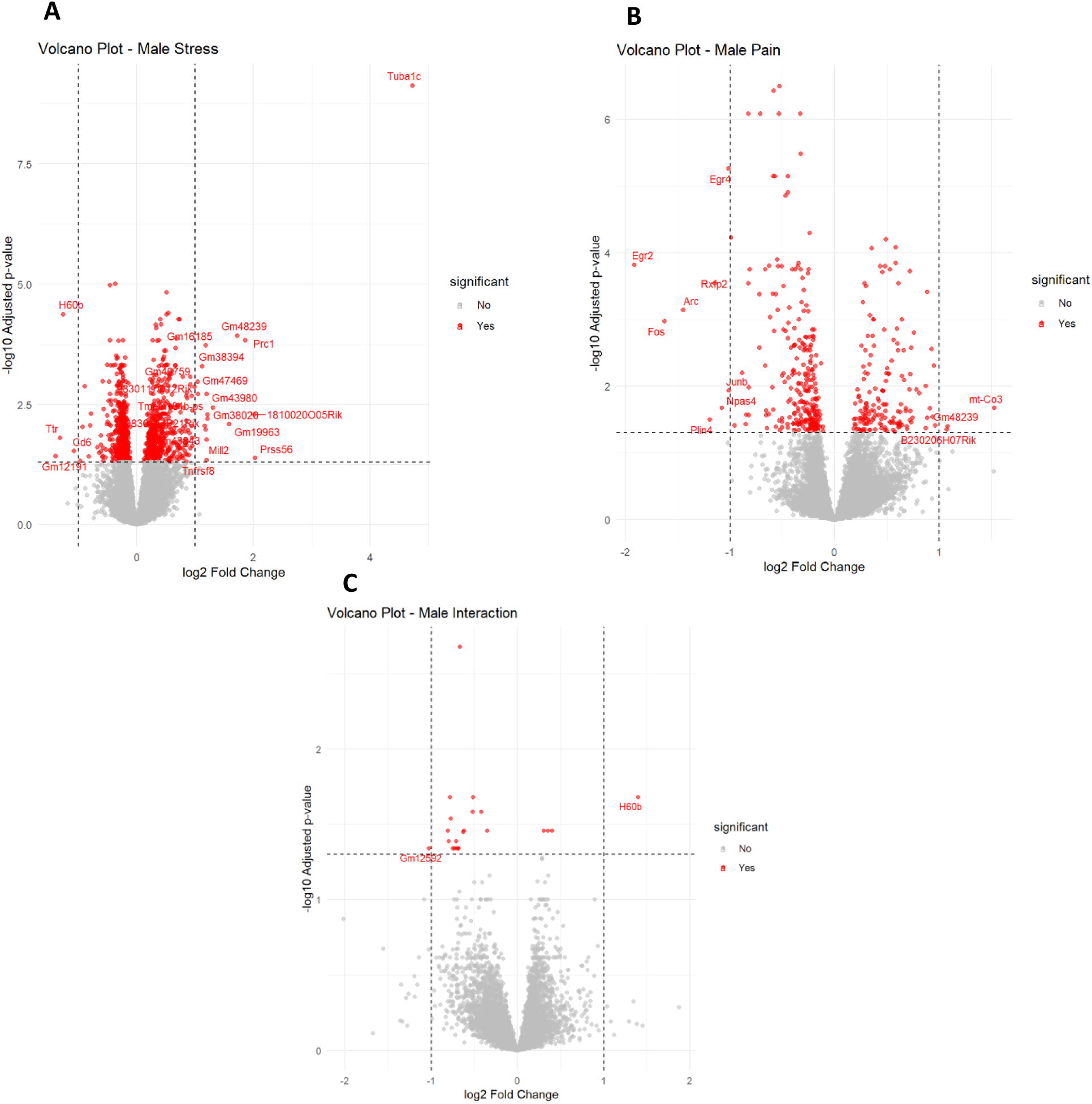
**Differential gene expression in the male frontal cortex following prenatal stress and chronic constriction injury (CCI)**. Differentially expressed genes identified by RNA sequencing. **(A)** Main effect of prenatal stress (prenatal stress vs sham). **(B)** Main effect of neuropathic pain (CCI vs sham). (**C**) Statistical interaction between prenatal stress and CCI, representing genes significantly regulated by the combined effect of both factors beyond their individual contributions. The x-axis represents log₂ fold change, and the y-axis shows the –log₁₀ adjusted *p*-value (FDR < 0.05). Genes passing significance thresholds are highlighted in red, with selected transcripts labeled.

### Functional Enrichment of Differentially Expressed Genes

To assess the biological significance of these transcriptional changes, we performed pathway enrichment analysis specifically on the interaction DEGs, as these genes are dependent on both PNS and CCI and are likely to be highly relevant to the observed increased pain sensitivity following PNS.

Enrichment analysis revealed that female interaction DEGs (Figure 4A) were associated with neuronal development, axonogenesis, and synaptic organization. In contrast, male interaction DEGs (Figure 4B) were enriched for ion transport, hypoxia response, and brain development, highlighting sex-dependent biological processes associated with transcriptional regulation dependent on both PNS and CCI.

**Figure 4.**
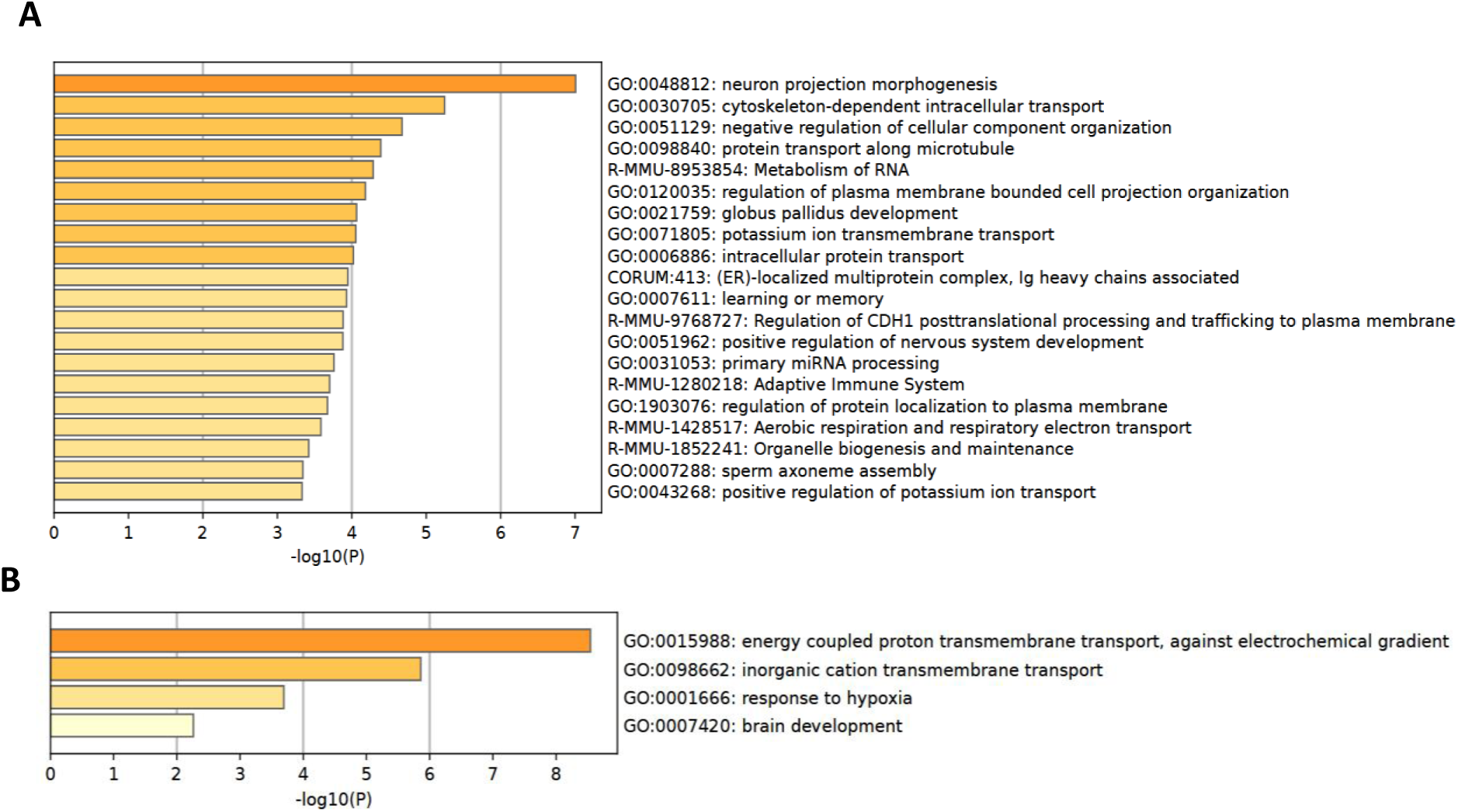
**Functional enrichment of differentially expressed genes (DEGs) in the frontal cortex of female and male offspring following PNS and CCI**. Metascape enrichment analysis of DEGs identified in the interaction analysis between PNS and CCI. Only interaction DEGs (p-adjusted < 0.05) were included. Bar plots display the top enriched GO biological processes ranked by –log₁₀ *p*-value in females (A) and males (B). Together, these findings demonstrate that PNS and CCI induce widespread and sex-specific transcriptomic alterations in the frontal cortex, converging on pathways central to neuronal signaling, immune regulation, and stress adaptation.

### DNA Methylation Alterations Following Prenatal Stress and CCI

To investigate whether PNS and CCI are associated with epigenetic alterations, we performed RRBS on frontal cortex samples. This analysis enabled the identification of DMRs linked to either PNS or CCI.

In females, prenatal stress resulted in 281 DMRs, while CCI produced 283 DMRs. In males, the number of methylation changes was substantially higher following prenatal stress, with 869 DMRs, whereas CCI was associated with 334 DMRs (Table 2). Since DMRs are short and highly localized it is unlikely interaction effect could be reliably detected, therefore for the RRBS analysis we focused on pair-wise comparisons.

**Table 2.** Number of differentially methylated regions (DMRs) in the frontal cortex following PNS and CCI. Counts of DMRs identified by RRBS in female and male offspring. Significant DMRs were defined as loci with pMWUq < 0.05.

| Effect | Female (DMRs) | Male (DMRs) |
| --- | --- | --- |
| Prenatal Stress (PNS) | 281 | 869 |
| CCI (Pain) | 283 | 334 |

To further visualize these methylation differences, volcano plots were generated for females (Figure 5A, C) and males (Figure 5B, D). In females, both PNS (Figure 5A) and CCI (Figure 5C) induced broad distributions of significant DMRs, with several loci exceeding 20% methylation change (ΔMethylation > 0.2) highlighted. In males, PNS (Figure 5B) displayed a strikingly larger spread of DMRs compared to CCI (Figure 5D), which produced a smaller and distinct set of DMRs.

**Figure 5.**
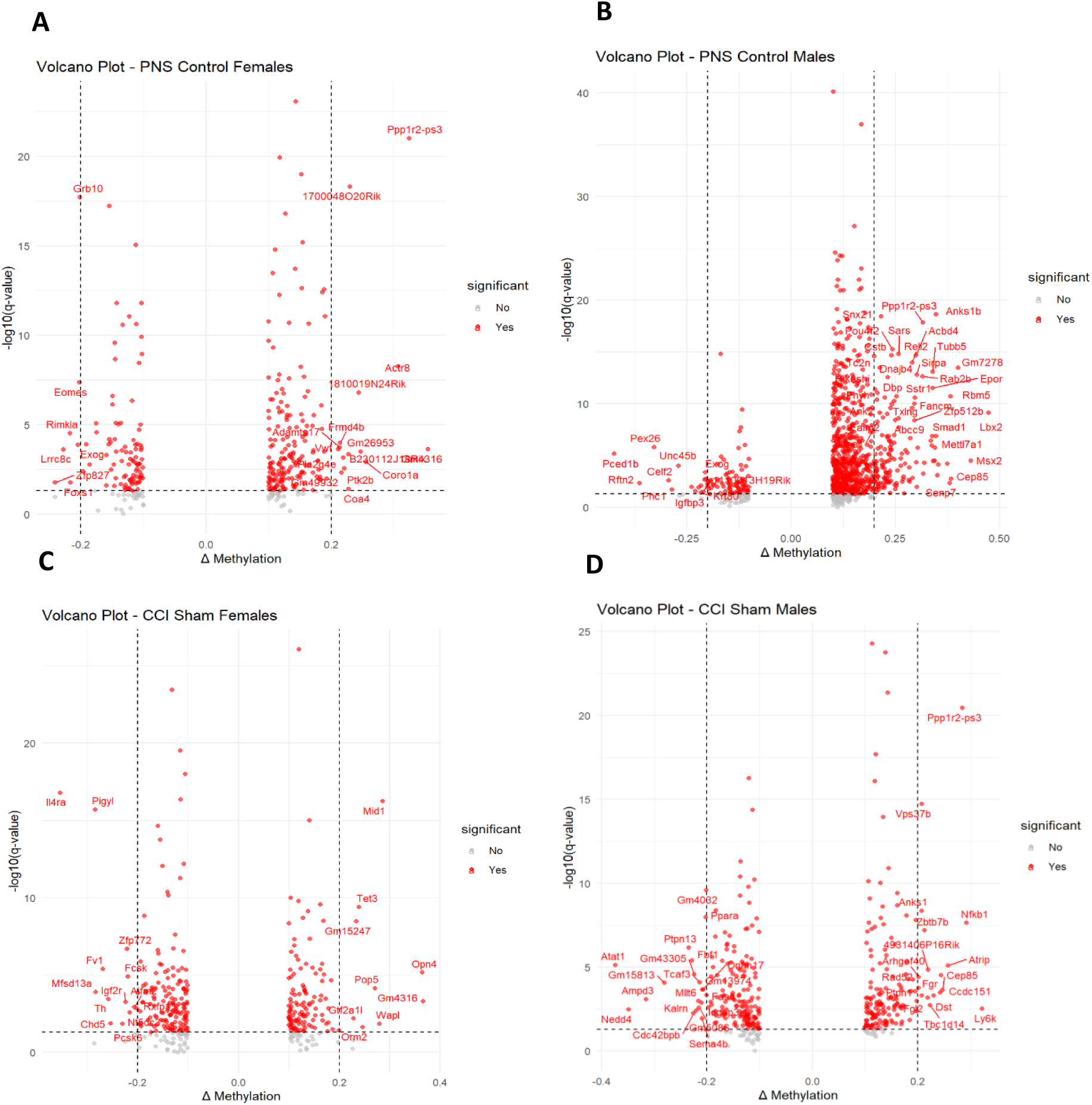
Differential DNA methylation in the frontal cortex of female and male offspring following prenatal stress and CCI. Volcano plots show DMRs identified by RRBS. (**A**) PNS females vs sham controls. (**B**) PNS males vs sham controls. (**C**) CCI females vs sham controls. (**D**) CCI males vs sham controls. The x-axis represents the change in methylation (ΔMethylation), and the y-axis represents the – log₁₀ *q*-value. Significant loci (pMWUq < 0.05) are highlighted in red, and regions with methylation changes greater than 20% (ΔMethylation > 0.2) are labeled. Numerous DNA regions not differentially methylated |<10%| sites were removed to allow for a clearer visualization.

### Functional Enrichment of DMR-Associated Genes

To determine the biological significance of these changes, we next performed pathway enrichment analysis of genes associated with significant DMRs. Following PNS (Figure 6), female DMR-associated genes (Figure 6A) were enriched for developmental and signaling processes, including cerebral cortex regionalization, axonogenesis, Wnt and Ephrin signaling, and neuron differentiation, together with synaptic regulation and cerebrospinal fluid circulation. In males (Figure 6B), enriched pathways were dominated by neuronal and synaptic functions such as synapse organization, regulation of synapse structure, neuron projection morphogenesis, and glutamate receptor signaling, alongside cytoskeletal remodeling, Rho GTPase and MAPK signaling, and developmental pathways including heart and skeletal system development.

**Figure 6.**
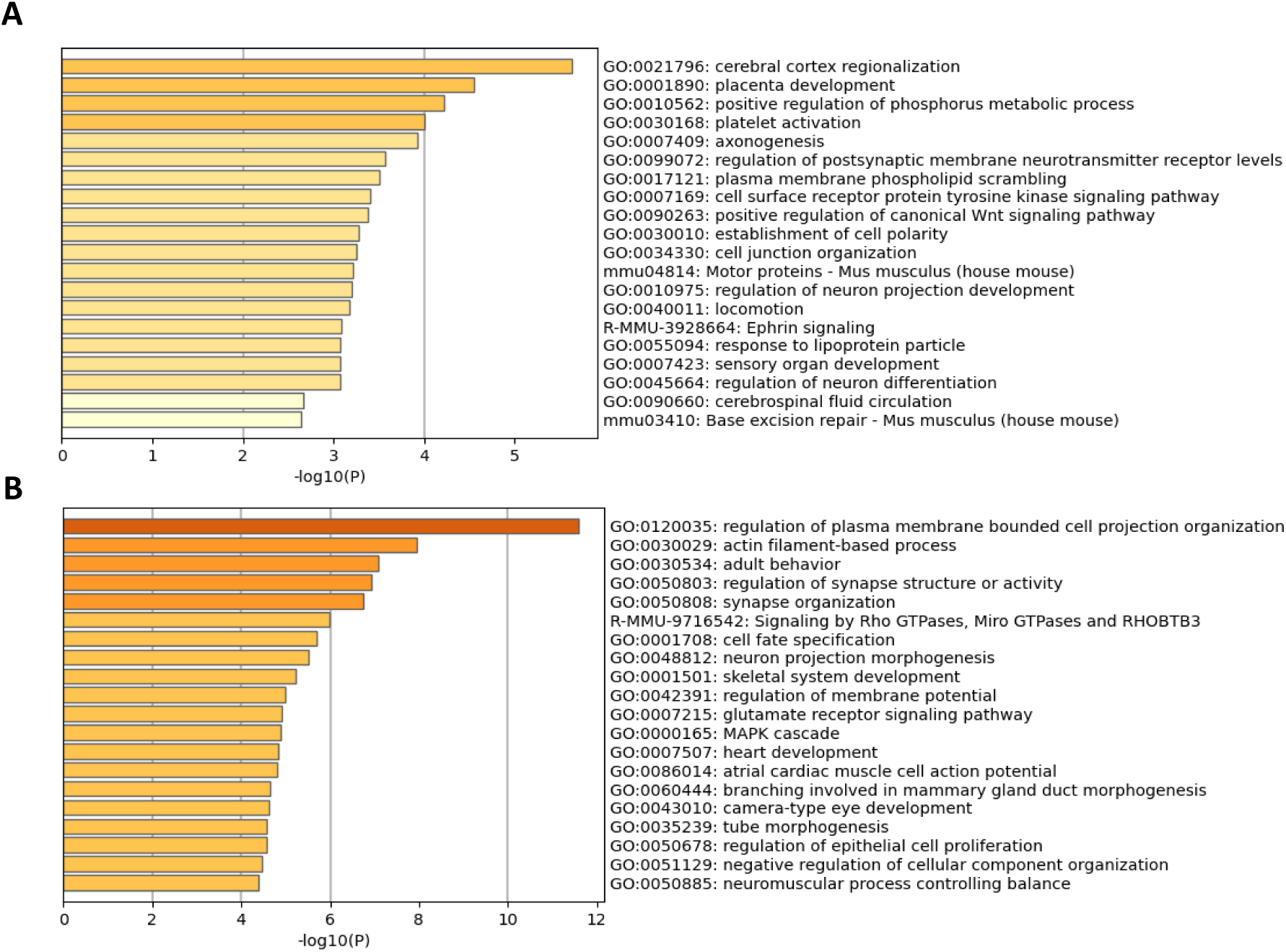
Functional enrichment of DMR-associated genes following PNS. Metascape analysis of genes associated with significant DMRs (pMWUq < 0.05) in the frontal cortex of (**A**) female and (**B**) male offspring exposed to PNS. Bar plots display the top enriched GO biological processes ranked by – log₁₀ *p*-value.

Following CCI (Figure 7), female DMR-associated genes showed enrichment for RNA splicing and synaptic protein regulation, cytoskeletal remodeling, carbohydrate metabolism, phosphatidylinositol signaling, and retinoic acid response. In males, enriched pathways included axon development, dendritic spine morphogenesis, receptor tyrosine kinase signaling, extracellular matrix organization, immune-related processes, and protein phosphorylation.

**Figure 7.**
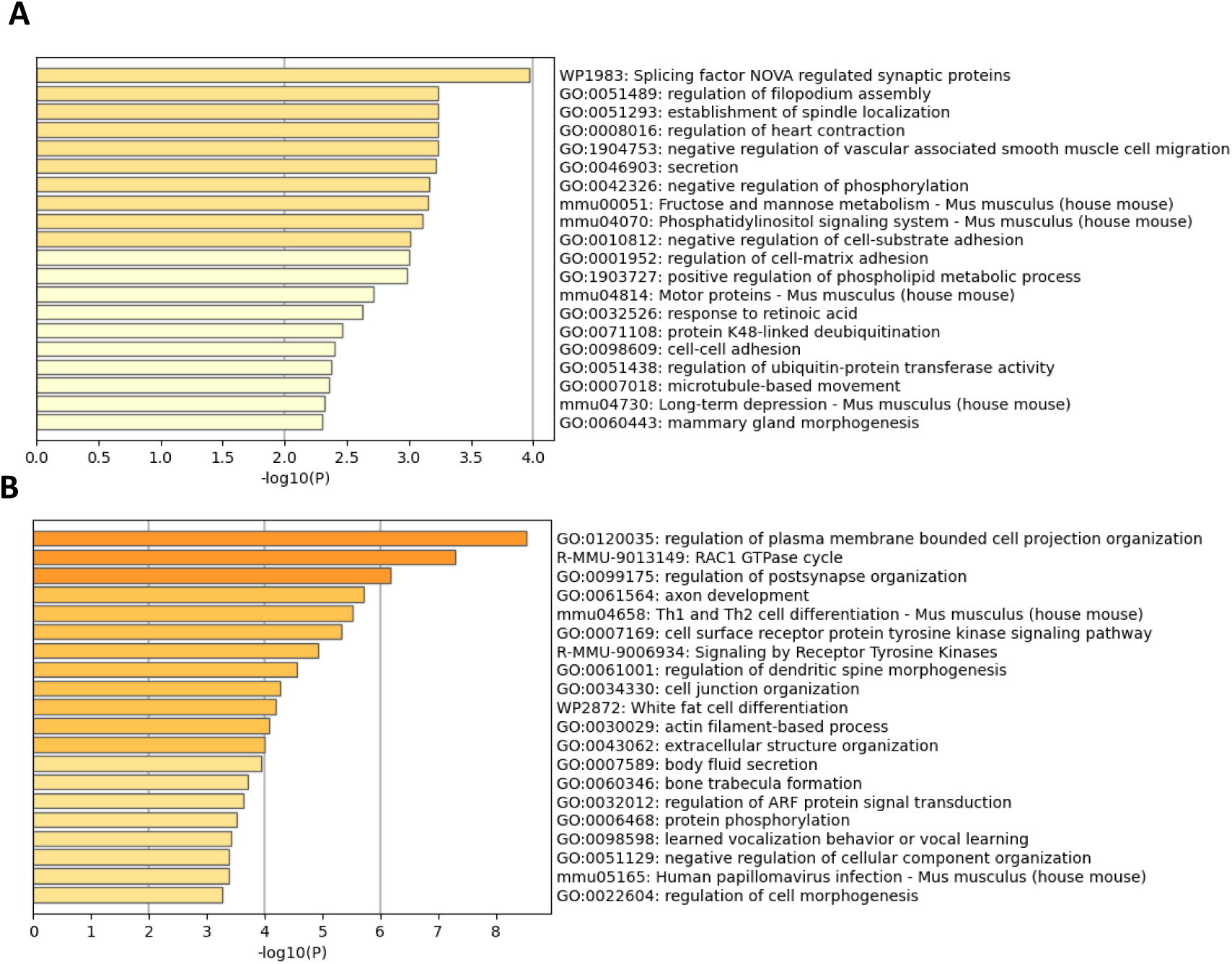
Functional enrichment of DMR-associated genes following CCI. Metascape analysis of genes associated with significant DMRs (pMWUq < 0.05) in the frontal cortex of (**A**) female and (**B**) male offspring following CCI. Bar plots display the top enriched GO biological processes ranked by –log₁₀ *p*-value.

Together, these results indicate that while both PNS and CCI induce DNA methylation changes in functionally relevant genes, the associated biological pathways differ between sexes and conditions, reflecting distinct epigenetic responses.

### Transcriptomic Effects of DMRs on Key Genetic Targets

To explore the transcriptional effects of DMR on key genes that were differentially methylated in several pair-wise RRBS analysis (Figure 8), we first overlaped the DMRs across conditions. We then prioritized eight targets with known roles in pain processing, neuronal development, or DNA methylation regulation (Tet3, Chd5, Pcsk6, Syngap1, Ptpn1, Cyfip2, Mbd2, Vgf) that were either present in two or more comparisons or were differentially expressed in our previous studies for quantitiative PCR analysis.

**Figure 8.**
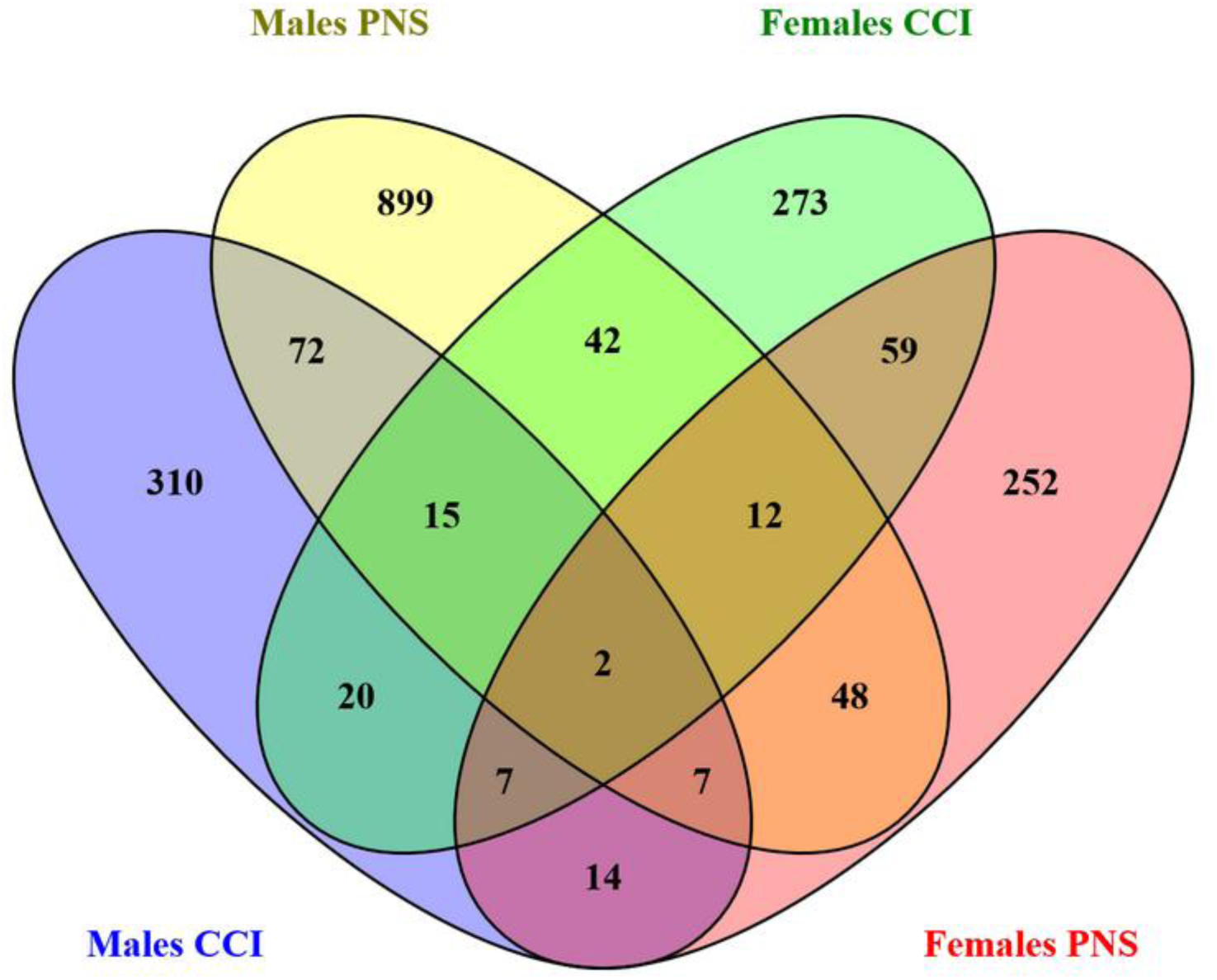
Venn diagram showing number of overlapped differentially methylated genes (FDR-correct p<0.05) between different comparisons

Using two-way ANOVA we found that CCI significantly reduced Tet3 levels in male (main effect (F (1, 27) = 13.87, P=0.0009)) but not in females (Figure 9A-B). Post hoc analysis found that Tet3 expression was signficantly lower in PNS-CCI males compared to control-sham and PNS-control.

**Figure 9.**
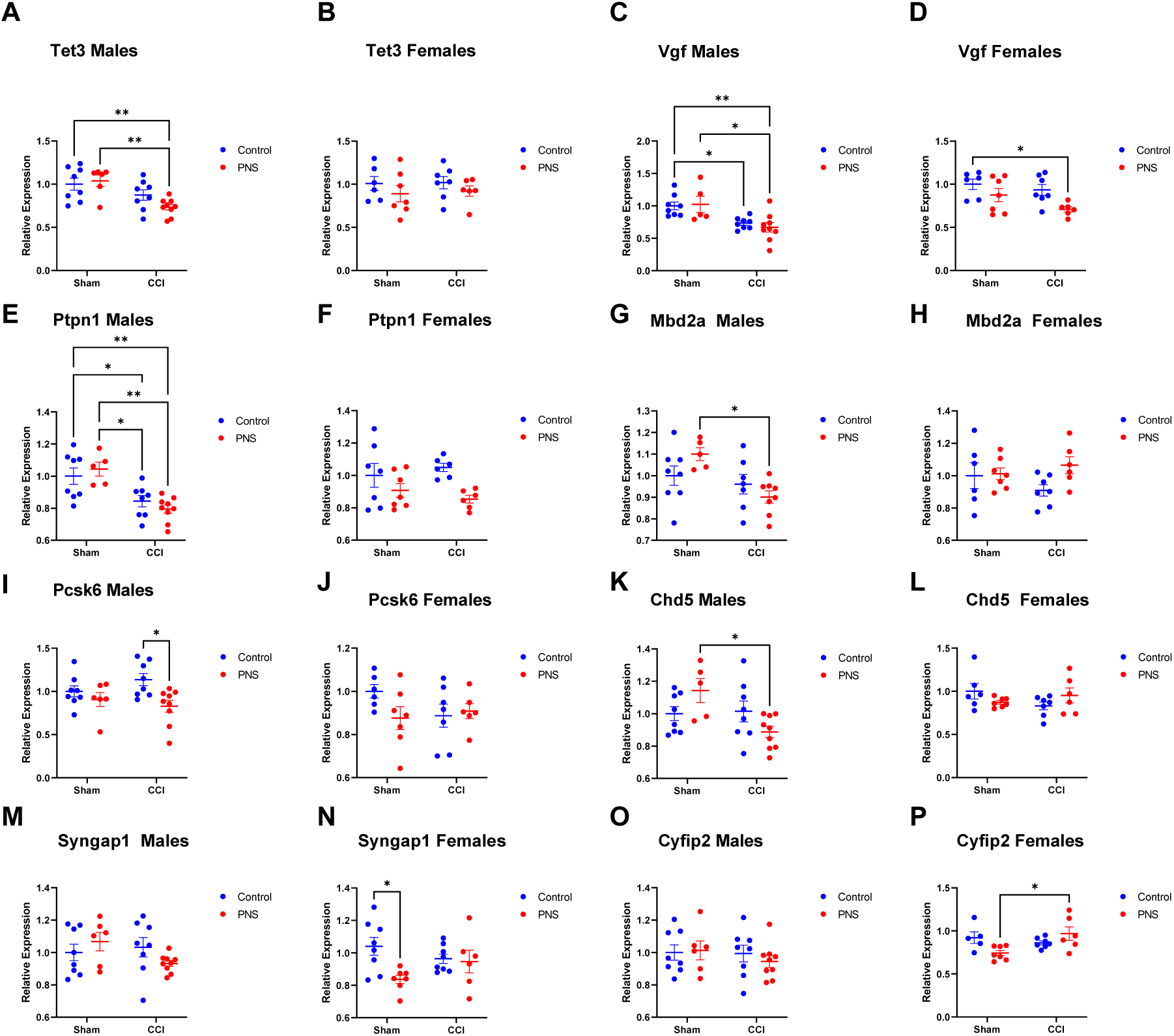
qPCR validation of mRNA expression of selected genes in the frontal cortex of male and female offspring. Relative mRNA levels of Tet3 **(A-B)**, Vgf **(C-D)**, Ptpn1 **(E-F)**, Mbd2a **(G-H)**, Pcsk6 **(I-J)**, Chd5 **(K-L)**, Syngap1 **(M-N)**, and Cyfip2 **(O-P)** were measured in Sham and CCI groups with or without PNS. Data points represent biological replicates; bars indicate mean ± SEM, Two-Way ANOVAs followed by post-hoc tests (*p < 0.05, **p < 0.01).

For Vgf we found that CCI reduced its expression in male (main effect F (1, 26) = 19.06, P=0.0002) while PNS reduced its expression in females (main effect F (1, 22) = 7.993, P=0.0098) (Figure 9C-D). Post hoc analysis in males found that PNS-CCI and control-CCI had significantly lower Vgf mRNA expression levels comapred to control-sham. In addition, PNS-CCI showed significanly reduced Vgf levels compared to PNS-sham. Post hoc analysis in females found PNS-CCI had significanly lower Vgf levels compared to control-sham.

Similarily, for Ptpn1, CCI reduced its expression in male (main effect F (1, 26 = 25.44, P<0.0001) while PNS reduced its expression in females (main effect F (1, 22) = 8.751, P=0.0073) (Figure 9E-F). The post-hoc analysis for male revealed that control-CCI and PNS-CCI mice had redcued Ptpn1 levels compared to both control-sham and PNS-sham. However, in females there were no significant changes in pair-wise comparisons.

For Mbd2a, there was a male-only effect with reduced levels in CCI (main effect F (1, 24) = 8.778, P=0.0068), and post hoc analysis found a PNS-CCI had significanly lower Mbd2a levels compared to PNS-sham. No significant effects were detected for females (Figure 9G-H).

Pcsk6 also showed a male-only effect with reduced levels in PNS mice (main effect F (1, 27) = 7.816, P=0.0094) and post hoc analysis found that PNS-CCI had lower Pcsk6 compared to control-CCI mice. No significant effects were detected for females (Figure 9I-J).

Male Chd5 levels were significantly lower in CCI (main effect F (1, 26) = 5.093, P=0.0327) and there was a PNS x CCI interaction in males (interaction F (1, 26) = 6.363, P=0.0181). The post hoc analysis found significantly reduced Chd5 levels in PNS-CCI males compared to PNS-sham, indicating that PNS is the main driver of reduced Chd5 in CCI mice. No significant effects were detected for females (Figure 9K-L).

In contrast to the aforementioned genes where most of the effects were seen in males, two genes were affected only in females. Specifically, PNS significantly reduced Syngap1 expression (main effect F (1, 25) = 5.576, P=0.0263), with post hoc analysis revealing that PNS-sham females had lower Syngap1 levels compared to control-sham. No significant effects were detected for males (Figure 9M-N).

For Cyfip2 there was a significant PNS x CCI interaction effect (interaction F (1, 21) = 7.875, P=0.0106) and post hoc analyis found signifcantly increased Cyfip2 levels in PNS-CCI females compared to PNS-sham suggesting that a dual effect of PNS and CCI is respnsible for upregulation of Cyfip2 in females. No significant effects were detected for males (Figure 9O-P).

## Discussion

Early pre- and post-natal stress, including PNS, is recognized as a major contributor to later life neuropsychiartic volunerability^50–52^ including vulnerability to chronic pain^17,53^. In addition, PNS exerts long-term effect on DNA methylation landscape in brain regions that mediate, at least partially, these vulnerabilities. Furthermore, we previously described that chronic pain induces persistant wide-spread changes on frontal cortex DNA methylation^36,37^. However, whether PNS effects on later life chronic pain hyper-sensitivity is associated with frontal cortex epigenetic alterations and downstream transcriptional changes, and whether there is a sex dicorcordance for these effects, is currently unkown.

Here, we demonstrated that both PNS and CCI induce widespread, long-term sex-specific alterations in gene-expression and DNA methylation. In addition, transcriptomic interaction effects suggest that PNS has a potentiating effect on CCI-induced alterations in some of the genes. These interaction effects were most pronounced in females with 660 DEGs than in males with only 22 DEGs. This might indicate that the combined effect of PNS on CCI is more robust for females.

In both females and males the interaction effects were strongly enriched for gene-pathways such as “neuron projection morphogenesis”, “protein transport along microtubule”, “brain development” indicating that neuronal morphology and organization was strongly affect by PNS and CCI. Other enriched pathways including “learning or memory” and “energy coupled proton transmembrane transport, against electrochemical gradients”, a key process that regulates membrane potential in neurons, suggesta the effect of PNS and CCI alters not only frontal cortex morphology but also cortical neuronal activity.

Our RRBS analysis showed genome-wide effects by PNS and CCI on the DNA methylome with stronger effects in males with a total of 1203 DMRs (869 for PNS and 334 for CCI) vs only 564 in females (281 PNS and 283 in CCI). These results are in line with our previous findings that spared nerve injury causes robust DNA methylation changes in the frontal cortex for up to 1 year post-injury^36,37^. Further, we previously showed for this cohort of mice that PNS and CCI alter the frontal cortex expression of epigenetic regulators including PNS-and CCI-induced reduction of Hdac1 in males and PNS and CCI-indcued elevation of Mbd2a in females^17^. These effects likely had downstream effects of both chromatin organization and DNA methylation maintanance leading to the observed genome-wide effects.

PNS-induced DNA methylation changes were enriched for gene pathways strongly related to neuronal development and morphology, neuronal activity and behaviors. Such pathways include: “cerebral cortex regionalization”, “axonogenesis”, “regulation of post synaptic membrane neurotransmitor receptor levels”, “locomotion”, “adult behavior” and “synapse organization” among others.

CCI-induced DNA methylation changes were enriched for gene pathways that include “splicing factor NOVA regulated synaptic plastisity”, “long term depression” and “axon development”, implying that genes differentially methylated by CCI are related to synaptic activity and neuronal structure.

Taken together, the methylomic and transcriptomic effects of both PNS and CCI converged into pathways highly relevant to altered brain structure and neuronal activity. These findings are in agreement with previous work showing altered sturcture, connectivity and function of the frontal cortex in chornic pain^54–57^ and in prenatal stress^17,58–60^. These changes, in turn, may be responsible for the long lasting effects of chronic pain and associated neuropsychiartic disoreders such as depression, anxiety and PTSD.

To further explore some of the main drivers of these effects we explored to which degree differentially methylated gene converge between pair-wise comparisons (Figure 8). We found several such genes that are also known for their role in pain processing (Vgf, Pcsk6, Ptpn1), neuronal development (Syngap1,Cyfip2) and epigenetic and chromatin regulation (Tet3, Chd5, Ptpn1, Mbd2). These genes were then examined for their mRNA expression levels by qPCR.

We found that both PNS and CCI reduced Pcsk6 in males but not in females with PNS-CCI group showing the lowest expression levels. In a human study, a genetic variation of this gene that encodes for the protease Proprotein Convertase Subtilisin/Kexin Type 6 protein (Pcsk6, also known as PACE4), was associated with protection against pain in knee osteoarthritis and this effect was successfully replicated in several independent^61^. Further, Psck6 knockout mice were significantly protected from pain across several mouse pain assays^61^. We found differentially methylated DMR annotated to Pcsk6 in all four comparisons (male and female PNS and CCI), suggesting this is a key gene in our model. It is possible that chronic pain induction and/or its persistence is regulated through Pcsk6 silencing and reduced pain resilience.

A similar expression pattern was found for Ptpn1 where PNS and CCI male mice both showed reduced expression. Ptpn1, also known as ptp1b, encodes for Protein Tyrosine Phosphatase 1B which is a regulator of neuroinflamatory singaling through Nf-Kb^62^. It was recently shown to be upregulated in spinal dorsal horn after spared nerve injury in rats^63^ and to regulate NMDA receptor-mediated peripheral inflammation-induced pain sensitization^64^. The observed cortical down regulation of this gene in our data might be a part of a neuroprotective response aiming to limit prologned neuroinflamation.

Vgf is a neurosecratory protein. It is a neuropeptide precursor that is upregulated by nerve injury and inflammation, and multiple VGF-derived peptides are pronociceptive and contribute to chronic pain hyper-sensitivity^65–68^. We found that Vgf was reduced by PNS and CCI in males and females. The finding of reduced Vgf in PNS is in agreement with previous reports of reduced Vgf in humans with major depressive disorder and in animal models of depression^65,66^.However, less is known about the effect of pain on cortical Vgf and our finding of reduced expression are contrary to the increased Vgf expression reported in injured sensory neurons^68–70^.

Syngap1 (Synaptic Ras GTPase Activating Protein 1) is part of the NMDA complex, making it a critical gene for neurodevelopment and neuronal function^71^. We found that PNS reduced Syngap1 expression in females, this might be a homeostatic response to PNS-induced stress in adulthood as lower Syngap1 levels, such as in Syngap1 heterozygote mice, are linked to reduced anxiety-like behaviors^72^.

Also Cyfip2 (Cytoplasmic FMR1 Interacting Protein 2), a protein involved in several processess including regulation of synaptic density and presynaptic function^73,74^ showed female-specific effects. There was a PNS x CCI interaction and PNS-CCI females showed increased Cyfip2 levels. This may indicate alteration in neurite outgrowth and synaptic function in this group.

Chd5 (Chromodomain Helicase DNA Binding Protein 5) is a neuron-specific helicase that is important for neurodevelopment and chromatin remodeling^75,76^, was significantly affected by both factors. Our analysis found a significant main effect of CCI causing a general reduction of Chd5 levels in male mice. Furthermore, we detected a significant PNS x CCI interaction, indicating that the effect of peripheral injury is not independent of prenatal history. The subsequent post hoc analysis confirmed that the most profound and significant transcriptional suppression of Chd5 occurs in the PNS-CCI group compared to the PNS-sham group. This specific finding suggests that prenatal stress sensitizes the male brain to subsequent injury.Tet3 (Tet Methylcytosine Dioxygenase 3) is an enzyme that convertes methylated cytosines into hydromethylated cytosines to promote de-methylation. Mbd2 (Methyl-CpG Binding Domain Protein 2) is a methylated-DNA “reader” that binds methylated DNA and stabilize DNA methylation. Both genes regulate brain function and behavior^40,77–80^. Tet3 and Mbd2 both showed reduced expression after CCI, when PNS-CCI resulted in even lower expression levels suggesting an additive or synergistic effect of PNS and CCI on their expression. MBD and TET proteins were reported to have either antagonistic effects on DNA methylation levels^81^ or potentially cooperative effects in some cases^82,83^. The similar expression patterns in our data suggest a transcriptional response that might be responsible for the widespread changes in the DNA methylation landscape.

Taken together, the effects of PNS and CCI on cortical DNA methylation and gene-expression show long lasting consequences of these insults, and thier combination, on brain function. Further, the highlighted key genes we tested were affected by prenatal stress and/or chronic pain in a sex-dependent manner. Future research is needed to explore the role of each of these and other genes in the molecular and behavioral effects of prenatal stress and its effects on chronic pain hyper-sensitivity in adulthood.

## Declaration of competing interest

None.

## Acknowledgements

This research was supported by a grant from the U.S.-Israel Binational Science Foundation (BSF) grant number #2021123 to EL and LSS. The authors thank Allison Maclean, Iabes Vilone, Peter Lee, The University of Minnesota Genomics Center and Aviad Sivan for technical support.

## Notes

### Competing Interest Statement

The authors have declared no competing interest.

## References

1. Institute of Medicine (US) Committee on Advancing Pain Research, Care, andEducation. *Relieving Pain in America: A Blueprint for Transforming Prevention, Care, Education, and Research*. (National Academies Press (US), Washington (DC), 2011).

2. Dahlhamer, J. et al. Prevalence of chronic pain and high-impact chronic pain among adults—United States, 2016. MMWR Morb. Mortal. Wkly. Rep. 67, 1001–1006 (2018).

3. Seminowicz, D. A. et al. Cognitive-behavioral therapy increases prefrontal cortex gray matter in patients with chronic pain. J. Pain 14, 1573–1584 (2013).

4. Thompson, J. M. & Neugebauer, V. Cortico-limbic pain mechanisms. Neurosci. Lett. 702, 15–23 (2019).

5. Baliki, M. N. et al. Corticostriatal functional connectivity predicts transition to chronic back pain. Nat. Neurosci. 15, 1117–1119 (2012).

6. Seminowicz, D. A. et al. Effective treatment of chronic low back pain in humans reverses abnormal brain anatomy and function. J. Neurosci. 31, 7540–7550 (2011).

7. Bilbao, A. et al. Longitudinal structural and functional brain network alterations in a mouse model of neuropathic pain. Neuroscience 387, 104–115 (2018).

8. Chan, J. C., Nugent, B. M. & Bale, T. L. Parental advisory: maternal and paternal stress can impact offspring neurodevelopment. Biol. Psychiatry 83, 886–894 (2018).

9. Cao-Lei, L. et al. DNA methylation signatures triggered by prenatal maternal stress exposure to a natural disaster: Project Ice Storm. PLoS One 9, e107653 (2014).

10. Nemoda, Z. & Szyf, M. Epigenetic alterations and prenatal maternal depression. Birth Defects Res. 109, 888–897 (2017).

11. Bussières, A. et al. Adverse childhood experience is associated with an increased risk of reporting chronic pain in adulthood: a stystematic review and meta-analysis. Eur. J. Psychotraumatology 14, 2284025 (2023).

12. Jones, G. T., Power, C. & Macfarlane, G. J. Adverse events in childhood and chronic widespread pain in adult life: results from the 1958 British Birth Cohort Study. Pain 143, 92–96 (2009).

13. You, D. S., Albu, S., Lisenbardt, H. & Meagher, M. W. Cumulative childhood adversity as a risk factor for common chronic pain conditions in young adults. Pain Med. 20, 486–494 (2019).

14. Macedo, B. B. D., von Werne Baes, C., Menezes, I. C. & Juruena, M. F. Child abuse and neglect as risk factors for comorbidity between depression and chronic pain in adulthood. J. Nerv. Ment. Dis. 207, 538–545 (2019).

15. Fumagalli, F., Molteni, R., Racagni, G. & Riva, M. A. Stress during development: impact on neuroplasticity and relevance to psychopathology. Prog. Neurobiol. 81, 197–217 (2007).

16. Nishinaka, T., Nakamoto, K. & Tokuyama, S. Enhancement of nerve-injury-induced thermal and mechanical hypersensitivity in adult male and female mice following early life stress. Life Sci. 121, 28–34 (2015).

17. Grégoire, S., Jang, S. H., Szyf, M. & Stone, L. S. Prenatal maternal stress is associated with increased sensitivity to neuropathic pain and sex-specific changes in supraspinal mRNA expression of epigenetic- and stress-related genes in adulthood. Behav. Brain Res. 380, 112396 (2020).

18. Moore, L. D., Le, T. & Fan, G. DNA methylation and its basic function. Neuropsychopharmacology 38, 23–38 (2013).

19. Weaver, I. C. G. et al. Epigenetic programming by maternal behavior. Nat. Neurosci. 7, 847–854 (2004).

20. McGowan, P. O. et al. Epigenetic regulation of the glucocorticoid receptor in human brain associates with childhood abuse. Nat. Neurosci. 12, 342–348 (2009).

21. Tajerian, M. et al. Peripheral nerve injury is associated with chronic, reversible changes in global DNA methylation in the mouse prefrontal cortex. PLoS One 8, e55259 (2013).

22. Alvarado, S. et al. Peripheral nerve injury is accompanied by chronic transcriptome-wide changes in the mouse prefrontal cortex. Mol. Pain 9, 21 (2013).

23. Massart, R. et al. Overlapping signatures of chronic pain in the DNA methylation landscape of prefrontal cortex and peripheral T cells. Sci. Rep. 6, 19615 (2016).

24. Chao, Y.-C. et al. Demethylation regulation of BDNF gene expression in dorsal root ganglion neurons is implicated in opioid-induced pain hypersensitivity in rats. Neurochem. Int. 97, 91–98 (2016).

25. Flavell, S. W. et al. Activity-dependent regulation of MEF2 transcription factors suppresses excitatory synapse number. Science 311, 1008–1012 (2006).

26. Ramamoorthi, K. et al. Npas4 regulates a transcriptional program in CA3 required for contextual memory formation. Science 334, 1669–1675 (2011).

27. Sun, L. et al. Contribution of DNMT1 to Neuropathic Pain Genesis Partially through Epigenetically Repressing Kcna2 in Primary Afferent Neurons. J. Neurosci. Off. J. Soc. Neurosci. 39, 6595–6607 (2019).

28. Ding, X. et al. DNMT1 Mediates Chronic Pain-Related Depression by Inhibiting GABAergic Neuronal Activation in the Central Amygdala. Biol. Psychiatry 94, 672–684 (2023).

29. Antunes, C., Sousa, N., Pinto, L. & Marques, C. J. TET enzymes in neurophysiology and brain function. Neurosci. Biobehav. Rev. 102, 337–344 (2019).

30. Pan, Z. et al. Hydroxymethylation of microRNA-365-3p Regulates Nociceptive Behaviors via Kcnh2. J. Neurosci. Off. J. Soc. Neurosci. 36, 2769–2781 (2016).

31. Shao, C. et al. DNMT3a methylation in neuropathic pain. J. Pain Res. 10, 2253–2262 (2017).

32. Sang, K. et al. Plastic change of prefrontal cortex mediates anxiety-like behaviors associated with chronic pain in neuropathic rats. Mol. Pain 14, 1744806918783931 (2018).

33. Liang, H.-Y. et al. nNOS-expressing neurons in the vmPFC transform pPVT-derived chronic pain signals into anxiety behaviors. Nat. Commun. 11, 2501 (2020).

34. Yin, J.-B. et al. dmPFC-vlPAG projection neurons contribute to pain threshold maintenance and antianxiety behaviors. J. Clin. Invest. 130, 6555–6570 (2020).

35. Zhuo, M. Long-term cortical synaptic changes contribute to chronic pain and emotional disorders. Neurosci. Lett. 702, 66–70 (2019).

36. Topham, L. et al. The transition from acute to chronic pain: dynamic epigenetic reprogramming of the mouse prefrontal cortex up to 1 year after nerve injury. Pain 161, 2394–2409 (2020).

37. Topham, L. et al. The methyl donor S-adenosyl methionine reverses the DNA methylation signature of chronic neuropathic pain in mouse frontal cortex. Pain Rep. 6, e944 (2021).

38. Zimmermann, M. Ethical considerations in relation to pain in animal experimentation. Acta Physiol. Scand. Suppl. 554, 221–233 (1986).

39. Zimmermann, M. Ethical guidelines for investigations of experimental pain in conscious animals. Pain 16, 109–110 (1983).

40. Lax, E. et al. Methyl-CpG binding domain 2 (Mbd2) is an epigenetic regulator of autism-risk genes and cognition. Transl. Psychiatry 13, 259 (2023).

41. Bennett, G. J. & Xie, Y.-K. A peripheral mononeuropathy in rat that produces disorders of pain sensation like those seen in man. Pain 33, 87–107 (1988).

42. Chaplan, S. R., Bach, F. W., Pogrel, J. W., Chung, J. M. & Yaksh, T. L. Quantitative assessment of tactile allodynia in the rat paw. J. Neurosci. Methods 53, 55–63 (1994).

43. Yoon, C., Wook, Y. Y., Sik, N. H., Ho, K. S. & Mo, C. J. Behavioral signs of ongoing pain and cold allodynia in a rat model of neuropathic pain. Pain 59, 369–376 (1994).

44. Dobin, A. et al. STAR: ultrafast universal RNA-seq aligner. Bioinformatics 29, 15–21 (2013).

45. Love, M. I., Huber, W. & Anders, S. Moderated estimation of fold change and dispersion for RNA-seq data with DESeq2. Genome Biol. 15, 550 (2014).

46. Krueger, F. & Andrews, S. R. Bismark: a flexible aligner and methylation caller for Bisulfite-Seq applications. Bioinforma. Oxf. Engl. 27, 1571–1572 (2011).

47. Akalin, A. et al. methylKit: a comprehensive R package for the analysis of genome-wide DNA methylation profiles. Genome Biol. 13, R87 (2012).

48. Love, M. I., Huber, W. & Anders, S. Moderated estimation of fold change and dispersion for RNA-seq data with DESeq2. Genome Biol. 15, 550 (2014).

49. Zhou, Y. et al. Metascape provides a biologist-oriented resource for the analysis of systems-level datasets. Nat. Commun. 10, 1523 (2019).

50. Provençal, N. & Binder, E. B. The effects of early life stress on the epigenome: From the womb to adulthood and even before. Exp. Neurol. 268, 10–20 (2015).

51. Peña, C. J. Early-life stress sensitizes response to future stress: Evidence and mechanisms. Neurobiol. Stress 35, 100716 (2025).

52. Miguel, P. M., Pereira, L. O., Silveira, P. P. & Meaney, M. J. Early environmental influences on the development of children’s brain structure and function. Dev. Med. Child Neurol. 61, 1127–1133 (2019).

53. Burke, N. N., Finn, D. P., McGuire, B. E. & Roche, M. Psychological stress in early life as a predisposing factor for the development of chronic pain: Clinical and preclinical evidence and neurobiological mechanisms. J. Neurosci. Res. 95, 1257–1270 (2017).

54. Apkarian, A. V. et al. Chronic back pain is associated with decreased prefrontal and thalamic gray matter density. J. Neurosci. Off. J. Soc. Neurosci. 24, 10410–10415 (2004).

55. Baliki, M. N., Chang, P. C., Baria, A. T., Centeno, M. V. & Apkarian, A. V. Resting-sate functional reorganization of the rat limbic system following neuropathic injury. Sci. Rep. 4, 6186 (2014).

56. Mayr, A. et al. Patients with chronic pain exhibit individually unique cortical signatures of pain encoding. Hum. Brain Mapp. 43, 1676–1693 (2022).

57. Kummer, K. & Sheets, P. L. Targeting Prefrontal Cortex Dysfunction in Pain. J. Pharmacol. Exp. Ther. 389, 268–276 (2024).

58. Barros, V. G., Duhalde-Vega, M., Caltana, L., Brusco, A. & Antonelli, M. C. Astrocyte-neuron vulnerability to prenatal stress in the adult rat brain. J. Neurosci. Res. 83, 787–800 (2006).

59. Fleming, D. E., Anderson, R. H., Rhees, R. W., Kinghorn, E. & Bakaitis, J. Effects of prenatal stress on sexually dimorphic asymmetries in the cerebral cortex of the male rat. Brain Res. Bull. 16, 395–398 (1986).

60. Wu, Y., De Asis-Cruz, J. & Limperopoulos, C. Brain structural and functional outcomes in the offspring of women experiencing psychological distress during pregnancy. Mol. Psychiatry 29, 2223–2240 (2024).

61. Malfait, A.-M. et al. A role for PACE4 in osteoarthritis pain: evidence from human genetic association and null mutant phenotype. Ann. Rheum. Dis. 71, 1042–1048 (2012).

62. Song, G. J. et al. A novel role for protein tyrosine phosphatase 1B as a positive regulator of neuroinflammation. J. Neuroinflammation 13, 86 (2016).

63. Jiao, B. et al. Protein tyrosine phosphatase 1B contributes to neuropathic pain by aggravating NF-κB and glial cells activation-mediated neuroinflammation via promoting endoplasmic reticulum stress. CNS Neurosci. Ther. 30, e14609 (2024).

64. Wu, S.-J. et al. Spinal PTP1B Regulated NMDA Receptor-mediated Nociceptive Transmission and Peripheral Inflammation-induced Pain Sensitization. Mol. Neurobiol. 62, 3781–3793 (2025).

65. Thakker-Varia, S. et al. The neuropeptide VGF produces antidepressant-like behavioral effects and enhances proliferation in the hippocampus. J. Neurosci. Off. J. Soc. Neurosci. 27, 12156–12167 (2007).

66. Jiang, C. et al. VGF and its C-terminal peptide TLQP-62 in ventromedial prefrontal cortex regulate depression-related behaviors and the response to ketamine. Neuropsychopharmacol. Off. Publ. Am. Coll. Neuropsychopharmacol. 44, 971–981 (2019).

67. Soliman, N., Okuse, K. & Rice, A. S. C. VGF: a biomarker and potential target for the treatment of neuropathic pain? Pain Rep. 4, e786 (2019).

68. Skorput, A. G. J. et al. Involvement of the VGF-derived peptide TLQP-62 in nerve injury-induced hypersensitivity and spinal neuroplasticity. Pain 159, 1802–1813 (2018).

69. Fairbanks, C. A. et al. The VGF-derived peptide TLQP-21 contributes to inflammatory and nerve injury-induced hypersensitivity. Pain 155, 1229–1237 (2014).

70. Riedl, M. S. et al. Proteomic analysis uncovers novel actions of the neurosecretory protein VGF in nociceptive processing. J. Neurosci. Off. J. Soc. Neurosci. 29, 13377–13388 (2009).

71. Agarwal, M., Johnston, M. V. & Stafstrom, C. E. SYNGAP1 mutations: Clinical, genetic, and pathophysiological features. Int. J. Dev. Neurosci. Off. J. Int. Soc. Dev. Neurosci. 78, 65–76 (2019).

72. Muhia, M., Yee, B. K., Feldon, J., Markopoulos, F. & Knuesel, I. Disruption of hippocampus-regulated behavioural and cognitive processes by heterozygous constitutive deletion of SynGAP. Eur. J. Neurosci. 31, 529–543 (2010).

73. Kim, Y. et al. Cell-autonomous reduction of CYFIP2 changes dendrite length, dendritic protrusion morphology, and inhibitory synapse density in the hippocampal CA1 pyramidal neurons of 17-month-old mice. Anim. Cells Syst. 28, 294–302 (2024).

74. Kim, G. H. et al. Altered presynaptic function and number of mitochondria in the medial prefrontal cortex of adult Cyfip2 heterozygous mice. Mol. Brain 13, 123 (2020).

75. Parenti, I. et al. Missense and truncating variants in CHD5 in a dominant neurodevelopmental disorder with intellectual disability, behavioral disturbances, and epilepsy. Hum. Genet. 140, 1109–1120 (2021).

76. Shrestha, P. et al. Chd5 Regulates the Transcription Factor Six3 to Promote Neuronal Differentiation. Stem Cells Dayt. Ohio 41, 242–251 (2023).

77. Antunes, C. et al. Tet3 ablation in adult brain neurons increases anxiety-like behavior and regulates cognitive function in mice. Mol. Psychiatry 26, 1445–1457 (2021).

78. Xie, D. et al. TET3 epigenetically controls feeding and stress response behaviors via AGRP neurons. J. Clin. Invest. 132, e162365 (2022).

79. Hendrich, B., Guy, J., Ramsahoye, B., Wilson, V. A. & Bird, A. Closely related proteins MBD2 and MBD3 play distinctive but interacting roles in mouse development. Genes Dev. 15, 710–723 (2001).

80. Wood, K. H. et al. Tagging methyl-CpG-binding domain proteins reveals different spatiotemporal expression and supports distinct functions. Epigenomics 8, 455–473 (2016).

81. Ludwig, A. K. et al. Binding of MBD proteins to DNA blocks Tet1 function thereby modulating transcriptional noise. Nucleic Acids Res. 45, 2438–2457 (2017).

82. Peng, L. et al. MBD3L2 promotes Tet2 enzymatic activity for mediating 5-methylcytosine oxidation. J. Cell Sci. 129, 1059–1071 (2016).

83. Wang, L. et al. Mbd2 promotes foxp3 demethylation and T-regulatory-cell function. Mol. Cell. Biol. 33, 4106–4115 (2013).

